# PathoPhenoDB: linking human pathogens to their disease phenotypes in support of infectious disease research

**DOI:** 10.1101/489971

**Authors:** Şenay Kafkas, Marwa Abdelhakim, Yasmeen Hashish, Maxat Kulmanov, Marwa Abdellatif, Paul N Schofield, Robert Hoehndorf

## Abstract

Understanding the relationship between the pathophysiology of infectious disease, the biology of the causative agent and the development of therapeutic and diagnostic approaches is dependent on the synthesis of a wide range of types of information. Provision of a comprehensive and integrated disease phenotype knowledgebase has the potential to provide novel and orthogonal sources of information for the understanding of infectious agent pathogenesis, and support for research on disease mechanisms. We have developed PathoPhenoDB, a database containing pathogen-to-phenotype associations. PathoPhenoDB relies on manual curation of pathogen-disease relations, on ontology-based text mining as well as manual curation to associate phenotypes with infectious disease. Using Semantic Web technologies, PathoPhenoDB also links to knowledge about drug resistance mechanisms and drugs used in the treatment of infectious diseases. PathoPhenoDB is accessible at http://patho.phenomebrowser.net/, and the data is freely available through a public SPARQL endpoint.

## Background & Summary

The 2016 Global burden of disease study estimated that infectious diseases as part of the communicable, maternal, nutritional and neonatal complex (CMNN) accounted for 19% of global mortality in 2016, and constitute the second most important cause of deaths globally [28]. They remain the top cause of mortality in most of the developing countries, mainly in Africa, at 56% in 2015. The annual infectious disease mortality in the world is reported as 10,598 (per 100,000 people) by the World Health Organization (WHO). Lower respiratory tract infections are the most likely cause of mortality due to infectious disease, followed by diarrhoeal diseases, tuberculosis and HIV/AIDS which were responsible for 3.2 million, 1.4 million, 1.4 million and 1.1 million deaths respectively in 2015 alone [3]. Infectious diseases have highly significant economic impact through morbidity and mortality, especially for the developing countries [5]. They affect multiple components of human development including income, health, education and productivity through lost life years, and cause devastating consequences worldwide.

Infectious diseases are caused by a wide range of organisms (viruses, bacteria, fungi, worms, protozoa) that are generally considered as pathogens. Antimicrobial drugs are often the first line therapy for infectious diseases. However, drug resistance accumulates over time due to selection of genetic changes in pathogen populations when they are exposed to antimicrobial drugs (such as antibiotics, antifungals, antivirals, antimalarials, and antihelmintics). It now becomes crucial to develop strategies that can identify a pathogen rapidly and determine successful treatment options based on functional information in the pathogen relevant to drug resistance mechanisms.

While functional information about pathogens and their interactions with hosts is increasingly becoming available on a molecular level through large-scale studies [29], phenotypes observed in a patient are not only mediated through direct molecular interactions between a pathogen and host but also through the immune response and physiological and patho-physiological processes affecting the entire host organism. Phenotypes observed in a patient provide a readout for all these processes and generally provide a proxy for the mechanism through with pathogens elicit their signs and symptoms [13]. While there is a wide range of phenotypes that are shared across multiple infectious diseases as a result of common immune system processes and immune response to pathogens, certain host-pathogen interactions may result in specific phenotypes through which pathogens can be broadly distinguished.

Phenotype-based computational analysis methods can uncover molecular mechanisms in Mendelian diseases [30], and have been applied to the discovery of disease mechanisms from animal models [16] and to the investigation of drug mechanisms and drug repurposing [15]. In the area of infectious disease, similar methods may be applicable, mainly to investigate mechanisms of virulence and pathogenicity. Application of phenotype-based methods requires matching phenotypes observed in a particular physiological or pathological state with the phenotypes known to be associated with pathogens [16, 14], and use of this information to reveal molecular mechanisms. Currently, there is no comprehensive database of pathogen-to-phenotype associations that can be used for this purpose.

We have developed PathoPhenoDB, a database of pathogen-to-phenotype associations intended to support infectious disease research. PathoPhenoDB is a database which relies on pathogen–disease associations curated manually from public resources and the scientific literature. We further expanded the pathogen–disease associations by complementary textmined data [20]. PathoPhenoDB links pathogens to their phenotypes based on manually-curated and text-mined disease–phenotype associations. Furthermore, PathoPhenoDB links pathogens to drugs [23] that are known to treat infections by the pathogen, and further links pathogens to drug resistance genes and proteins [19] as well as to the drugs against which these genes or proteins convey resistance so that the information in PathoPhenoDB can be utilized directly for research on drug resistance mechanisms. PathoPhenoDB is freely available on http://patho.phenomebrowser.net, and the data can be obtained through a public SPARQL endpoint.

## Methods

### Data collection and integration

We developed PathoPhenoDB by considering the FAIR data principles [38]. We gathered a list of possible human pathogens for all infectious disease listed in the DO [22]. We identified possible pathogens for these diseases from public databases and scientific literature. In the cases where we could not find an exact match of a pathogen in the NCBI taxonomy, we mapped the pathogen to their parent class. For example, instead of assigning *Spirillium minus* to *Sodoku disease*, we assigned the higher taxon *Spirillium* to *Sodoku disease* due to *Spirillium minus* not being listed in the NCBI taxonomy.

We extracted disease phenotypes manually from DO and largely from PubMed abstracts by using ontology-based text mining [17], and we then linked the phenotypes to the pathogens. Briefly, we identified co-occurrences between disease names from DO and phenotype names from the Human Phenotype Ontology (HPO) [33] (downloaded on 14/May/2018) and the Mammalian Phenotype Ontology (MP) (downloaded on 14/May/2018) [9] in abstracts. We selected the significant disease–phenotype co-occurrences based on an ontology-based normalized point-wise mutual information [17]. To integrate the HPO and MP ontologies we use the PhenomeNET ontology [16].

We extracted drug information from the SIDER database [23] by resolving the cross-references between UMLS concept identifiers and DO identifiers. We gathered information about drug resistance from the Antibiotic Resistance Ontology (ARO) [19]. For this purpose, we matched the ARO accession and iterated through the ontology hierarchy using the subclass and *confers-resistance-to* relationships in ARO to retrieve the drugs. Using the Entrez API, we then identified the DNA accession that conveys drug resistance in the NCBI nucleotide database to retrieve the organism and its NCBI Taxonomy identifier. While, diseases are linked to pathogens, drugs, and phenotypes, the pathogens are linked to their drug resistance proteins, DNA accession and drugs to which they are resistant.

### Semantic similarity computation

We calculate the semantic similarities by using Resnik’s semantic similarity measure [32]. We rank pathogens based on their similarity scores to the query to find candidate pathogens. The Resnik semantic similarity measurement is formally defined as:

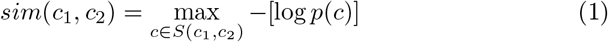

where *p*(*c*) is the probability of a pathogen being annotated with *c*. We used the Best-Match Average strategy to calculate the average of of all maximum similarities and compare two sets of phenotypes:

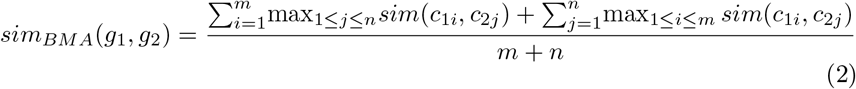

Additionally, we use the OPA2Vec framework [35] to generate ontology embeddings for the pathogens, and we use a t-SNE dimensionality reduction [26] to visualize the resulting embeddings and their relations.

### Implementation and availability

We developed the web application for our database using Python Django framework [1] for the backend and ReactJS [2] for the frontend services. We store only identifiers for the links in our database and retrieve all the annotations and additional data such as names from the AberOWL [34] ontology repository using its REST API. For this purpose, we created compressed versions of DO, PhenomeNET and NCBITAXON [10] ontologies.

PathoPhenoDB is available at http://patho.phenomebrowser.net and its content is accessible through a public SPARQL endpoint. The source code is available at https://github.com/bio-ontology-research-group/pathophenodb and every release of the data is deposited in a research data repository (Data Citation 1).

## Data records

PathoPhenoDB is a database of human pathogens, the diseases and phenotypes they elicit in human organisms, and information related to drug treatments and mechanisms of drug resistance. Specifically, PathoPhenoDB contains associations between pathogens and diseases, between pathogens and phenotypes, between drugs that are approved to treat particular pathogens, and it identifies genes or proteins within pathogens that can convey resistance to particular drugs. Figure 1 provides a high-level overview of the information in PathoPhenoDB. PathoPhenoDB is constructed through a combination of manual curation of scientific literature, text mining, information extraction, and data integration approaches using Semantic Web technologies.

**Figure 1:**
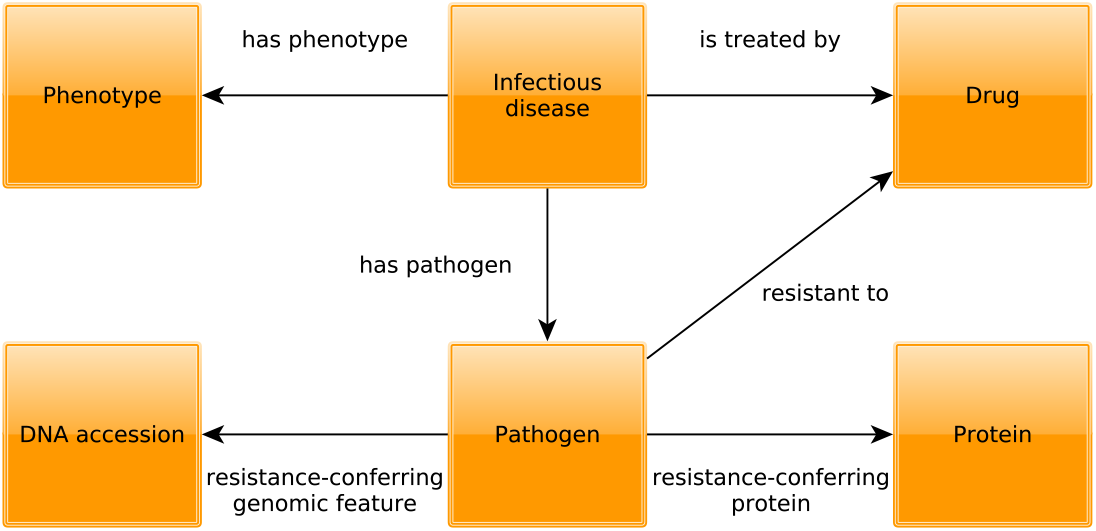
Schematic overview of the types of entities and their relations in PathoPhenoDB

In PathoPhenoDB, we consider a pathogen to be any kind of bacterium, virus, fungus, protozoan, parasite, or other type of organism that is known to be able to cause a disease or abnormal phenotype in humans. With this broad view of pathogens, PathoPhenoDB includes 32 types of parasitic insects, 115 fungi, 208 bacteria, 47 protozoa, 175 viruses, and 115 taxa of parasitic worms. Our database currently covers a total of 1,170 pathogen–disease associations between 508 infectious diseases and 692 taxa of pathogens. For a total of 130 infectious diseases and 399 pathogen–disease associations, we also include information about drugs that can treat the disease and pathogen. We further include information on known mechanisms of drug resistance for 30 pathogens representing 78 pathogen–disease associations. While PathoPhenoDB is largely based on manually curated information, we also extracted pathogen–disease associations from the biomedical literature [20] and use this information to enrich the content of our database. Statistics relevant to the textmined content is available from the web site http://patho.phenomebrowser.net.

We use the Human Disease Ontology (DO) [22] as a reference for infectious diseases in humans and base all our disease-related information on the DO. To associate pathogens with phenotypes, we follow a data integration approach and deductive inference where we utilize pathogen–disease associations to propagate phenotypes associated with infectious disease in DO [17] to the pathogens that cause the diseases. In PathoPhenoDB, we utilize 1,140 of 1,143 pathogen– disease associations to assign phenotypes for 476 (out of 508) infectious diseases to pathogens. For example, we use the phenotypes assigned to the DO class *Plasmodium malariae malaria* (`DOID:14324`), which includes phenotypes such as “episodic fever”, “hemolysis” and “anura”, and assign all phenotypes of this disease to *Plasmodium malariae* (`NCBITaxon:5858`) based on the association between *Plasmodium malariae* and *Plasmodium malariae malaria*.

As vocabulary for phenotypes we use a combination of the Human Phenotype Ontology (HPO) [33] and the Mammalian Phenotype Ontology (MP) [37]. While both ontologies are formally distinct and use different identifiers, they can be integrated and aligned through cross-species phenotype ontologies such as PhenomeNET [16] or UberPheno [25]. For our database, we use the PhenomeNET ontology as it has been applied in a variety of phenotype-driven studies of molecular mechanisms [4, 24]. The 692 pathogens in PathoPhenoDB are associated with 1,719 distinct phenotypes from HPO and 479 distinct phenotypes from MP. On average, each pathogen is directly associated with 20 phenotypes.

Using ontologies to represent phenotypes enables deductive inferences using the ontology axioms [27, 18]. To exploit these inferences, we represent the data in PathoPhenoDB using the RDF format (described in the methods section) and it is available for querying by using SPARQL endpoint from the web site. We use the Relation Ontology (RO) [36] and the Semanticscience Integrated Ontology [8] to represent the relations between the entities. For the relations in Figure 1, the `has_phenotype` relation corresponds to `RO_0002200, has_pathogen` to `RO:0002556` and the `is_treated_by` relation to `RO:0002302`.

As we gathered pathogen–disease and disease–phenotype associations using two different methods – text mining and manual curation – we use the `has_annotation` relation from SIO (`SIO:000255`) to reify annotation assertions; in particular, we generate annotation objects that consists of a relation, an annotation value of the relation, and an evidence code that represents the level of evidence for the annotation. For example, to represent the information that *Actinomadura madurae* (`NCBITaxon:1993`) may cause *Actinomycosis* (`DOID:8478`), as obtained from text mining, we generate a new annotation object consisting of the *pathogen of* (`RO:0002556`) relation to *Actinomycosis* (`DOID:8478`) and the *has evidence* (`RO:0002558`) relation to *manual assertion* (`ECO:0000203`) in the Evidence and Conclusion Ontology (ECO) [12]. In addition to reusing relations from established ontologies, we generate new relations (*resistant to*, *resistance-conferring protein*, *resistance-conferring genomic feature*) to capture information on drug resistance.

The data in PathoPhenoDB is updated when new data becomes available or issues with existing data are resolved. To report issues such as incorrect or missing associations, we maintain an issue tracker. Data is released in RDF format and every release of the data is deposited in the Zenodo data repository (Data Citation 1).

## Technical Validation

Figure 2 presents the search in PathoPhenoDB for a particular phenotype, *Brain atrophy*. The query retrieves all the infectious diseases associated with brain atrophy and their causative pathogens. Due to the use of inference and Semantic Web technologies, PathoPhenoDB can identify both direct and indirect associations between pathogens and phenotypes. We classify an association as direct if it is explicit asserted during the curation. We classify an association as indirect if it is inferred based on a direct association and application of inference over the subclass hierarchy of the ontologies. For example, PathoPhenoDB does not cover any infectious disease that is directly associated with *Toxocara* (`NCBITaxon:6264`). However, a query for *Toxocara* will return all the disease and phenotype associations linked to its subclasses, including *Toxocara canis* (`NCBITaxon:6265`) and *Toxocara canti* (`NCBITaxon:6266`) as indirect associations. Using this kind of inference, PathoPhenoDB can provide useful and relevant knowledge on the entities of interest while using the background knowledge contained in the class hierarchy of the ontologies.

**Figure 2:**
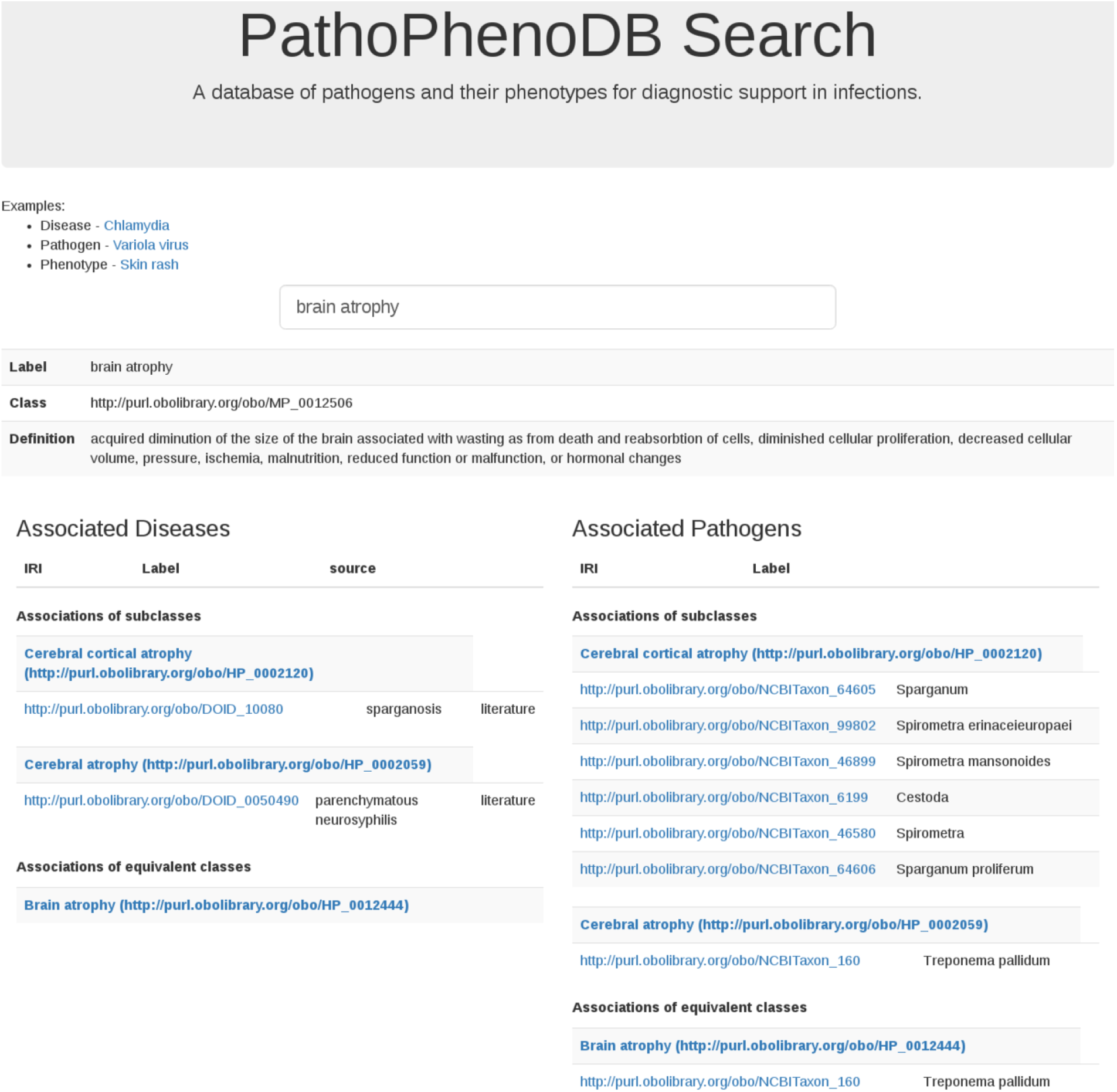
Search in PathoPhenoDB.

In addition to querying our database using Semantic Web technologies, the information in PathoPhenoDB can also be used to perform approximate queries using semantic similarity measures [32]. We test the similarity-based retrieval of pathogens by generating synthetic sets of phenotypes that consist of randomly chosen subsets of phenotypes associated with an infectious disease, and trying to identify the pathogen causing the disease based on semantic similarity over phenotype ontologies. Figure 3 shows the performance of recovering pathogens through semantic similarity when providing a varying number of symptoms. The model achieves a ROCAUC of over 86% using a single phenotype as query and does not significantly improve with more phenotypes added. We speculate that this is the result of pathogens falling in distinct phenotypic groups and that semantic similarity does not appropriately weight the distinguishing phenotypes within the group (because they are too general). We further use a data-driven semantic similarity measure based on ontology embeddings [35] to visualize the pathogens and their phenotypes in a 2-dimensional space using a t-SNE dimensionality reduction [26]; phenotypically related pathogens are closer together in this space. Figure 4 shows the resulting plot. We also make this figure available on our website to enable interactive exploration of pathogens based on their phenotype similarity.

**Figure 3:**
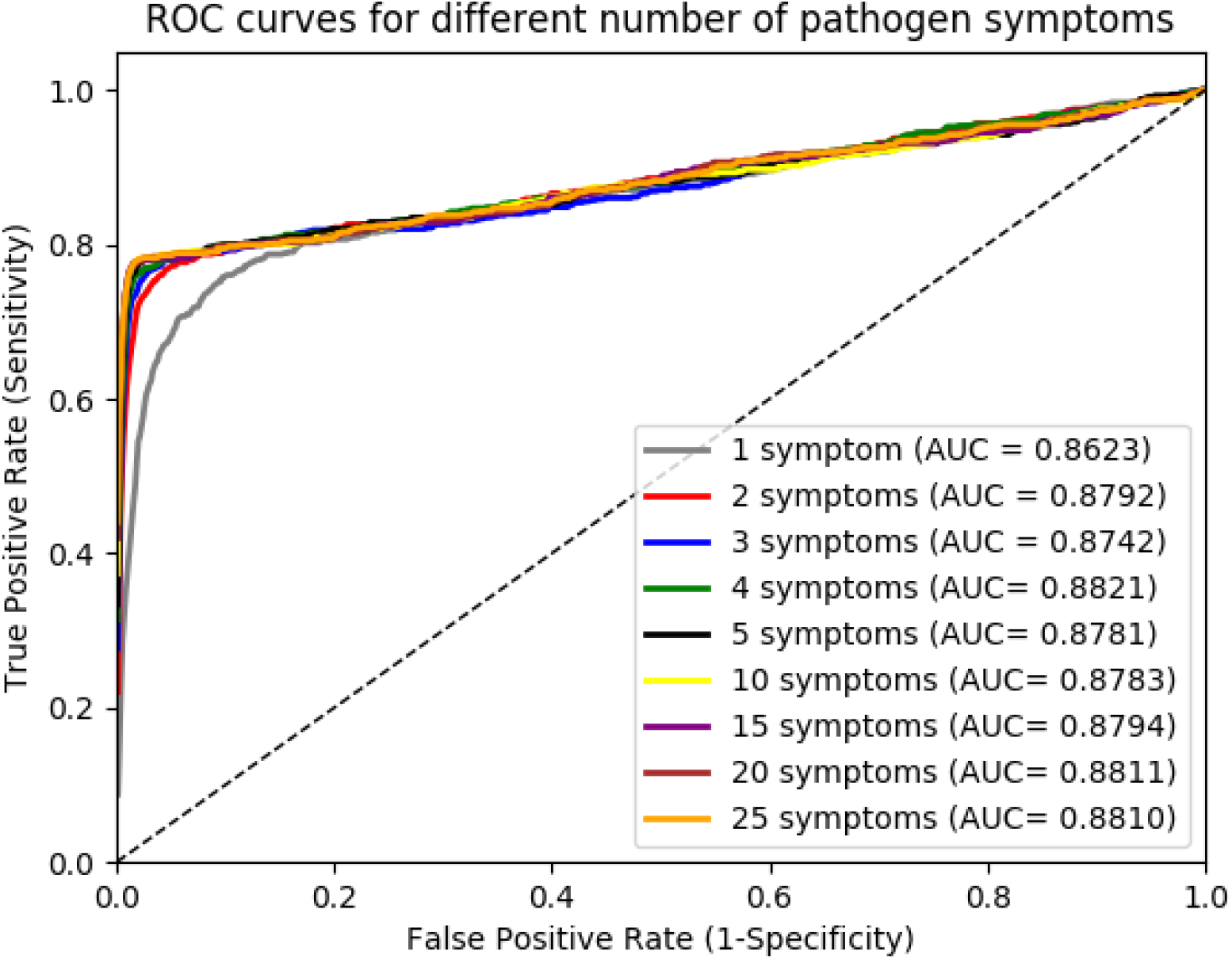
Pathogen recovery with different number of symptoms.

**Figure 4:**
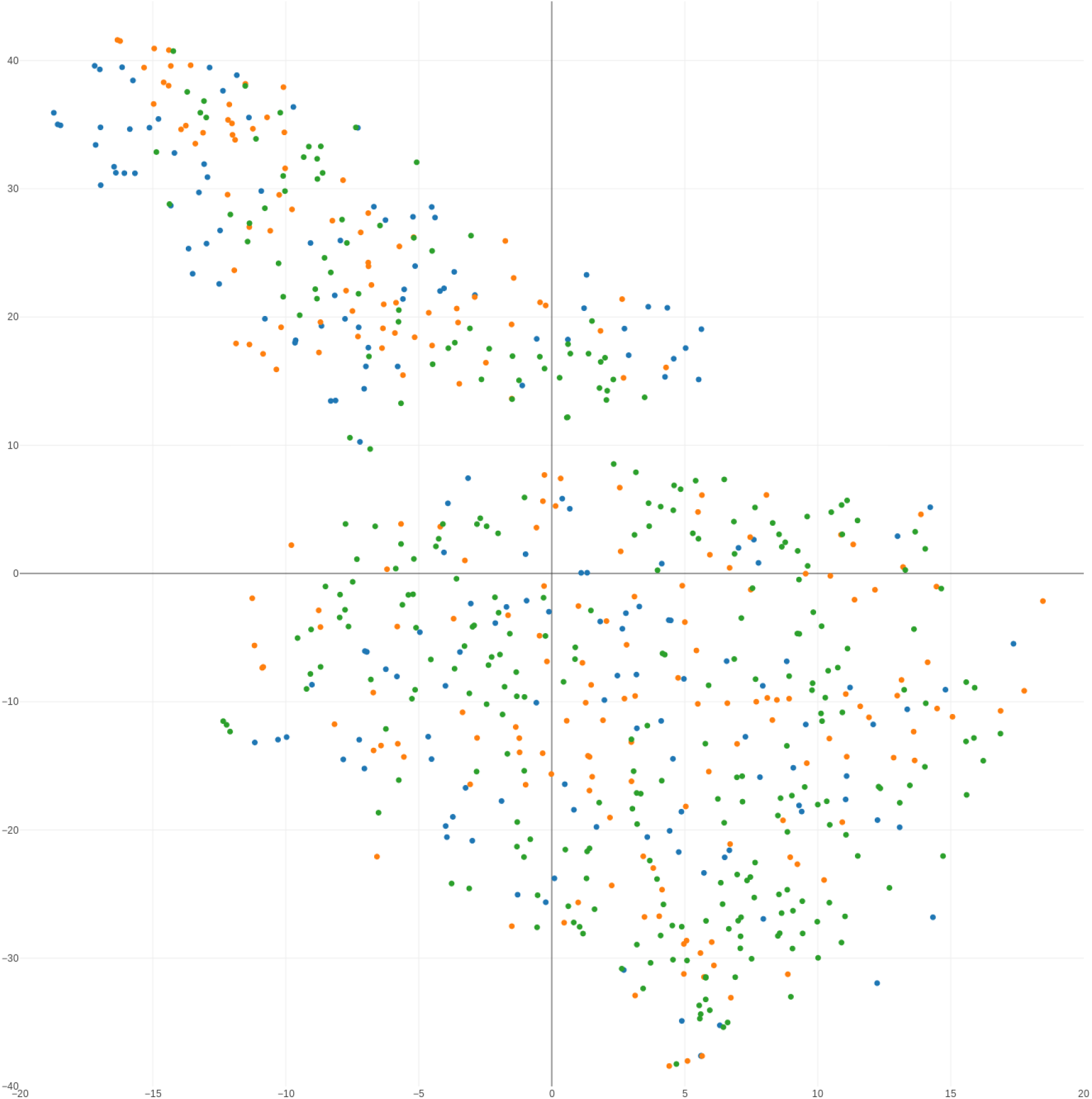
t-SNE plot of pathogens. Pathogens are represented using their ontology embeddings that have been generated using their associated phenotypes. Viruses are colored in blue, bacteria in orange, all other pathogens in green.

## Usage notes

Infectious disease research and diagnosis of infectious disease is rapidly changing with the application of sequencing technologies. Current routine clinical pathogen identification methods often do not identify the most effective and specific treatment options [6], or are not able to identify the causative pathogens rapidly. Recent achievements in next generation sequencing technologies (NGS) have led clinical microbiology to move in the direction of molecular diagnostic approaches [7]. NGS, in particular metagenomics and metatranscriptomics, can address the limitations of traditional microbial diagnostic methods by offering unbiased identification of organisms and can also be used to identify drug resistance and other functional information. Furthermore, metagenomics enables us to detect non-culturable organisms and multiple infections, and already shows great potential to be used in the rapid and accurate identification of pathogens [11, 31].

While NGS-based approaches have the potential to identify a wide range of pathogens in a single sequencing run, they may identify multiple different microorganisms that have the potential to cause infections. Identification of the causative pathogen among the set of pathogens identified through metagenomics approaches, is an additional challenge. In the future, matching the phenotypes observed in a patient to phenotype in PathoPhenoDB may provide additional features that can be combined with information from NGS to improve diagnosis and treatment of infectious disease.

As a more direct application of PathoPhenoDB, we envision its use in investigating molecular mechanisms underlying infectious diseases, specifically host-pathogen interactions. Phenotypes indirectly encode the molecular interactions between hosts and pathogens and therefore may be used to study the molecular basis of infectious disease.

## Acknowledgements

This work was supported by funding from King Abdullah University of Science and Technology (KAUST) Office of Sponsored Research (OSR) under Award No. URF/1/3454-01-01 and FCC/1/1976-08-01.

## Author contributions

Ş.K., Y.H. and M.A. contributed to the extraction and manual curation of the data. Ş.K. performed the experiments. Ş.K., R.H. and P.S. analyzed the results. M.K. and Marwa Abdellatif designed the web interface. Ş.K. drafted the manuscript. All authors revised, reviewed and approved the final version of the manuscript.

